# Targeting IL-1 Signaling to Improve Antidepressant Efficacy in a Rodent Model of Postpartum Depression

**DOI:** 10.64898/2026.11.26.769549

**Authors:** Romina Garcia de leon, Wansu Qiu, Phoenix Au-Yeung, Stephanie E Lieblich, Amanda B. Namchuk, Liisa A.M Galea

## Abstract

The postpartum is a period of heightened vulnerability to first-onset depression, with postpartum depression (PPD) affecting approximately 19% of birthing parents. Despite the widespread use of selective serotonin reuptake inhibitors (SSRIs) to treat PPD, remission rates remain low, and the mechanisms underlying limited antidepressant efficacy are poorly understood. Emerging preclinical evidence implicates hippocampal interleukin (IL)-1β signalling in SSRI efficacy during the postpartum. Here, we investigated whether IL-1 receptor antagonism facilitates the therapeutic effects of fluoxetine (FLX) in a preclinical model of PPD. High corticosterone (CORT) administration for 21 days postpartum disrupted maternal care, increased passive coping, and reduced indices of neuroplasticity, as evidenced by decreased hippocampal neurogenesis and increased numbers of perineuronal net+ neurons in the hippocampus and frontal cortex. Combined treatment with FLX and anakinra (KIN), an IL-1 receptor antagonist, reversed CORT-induced alterations in neuroplasticity, whereas neither treatment alone was effective. However, these neuroplasticity effects were dissociable from behavioural outcomes, as combined treatment failed to reverse increased passive coping in the forced swim test. These findings implicate IL-1 signalling as a potential constraint on antidepressant-induced neuroplasticity during the postpartum and identify IL-1 receptor antagonism as a promising adjunctive strategy to overcome limited neuroplastic responses to SSRIs in PPD.

## INTRODUCTION

Perinatal depression (PND) is a subtype of major depressive disorder (MDD) defined as depression during gestation or within 4 weeks postpartum [1]. However, there are distinct differences in biomarkers and risk factors based on timing of symptom onset (gestational versus postpartum) for PND, yet few studies distinguish between these periods [2,3]. First-time depression is more common during the postpartum [4–7] with the prevalence of postpartum depression (PPD) as high as 17-19% globally [8,9]. Given the increased risk for first onset of depression in the postpartum, it is important that we understand the underlying biology of postpartum depression onset as it relates to risk and therapeutic efficacy.

Dysregulation of the hypothalamic–pituitary–adrenal (HPA) axis is a feature of both major depressive disorder (MDD) and PPD. Notably, women with depression onset during the postpartum exhibit higher basal cortisol levels than those with depression onset during or before pregnancy [10], implicating elevated glucocorticoids in the pathophysiology of PPD. Consistent with these clinical findings, chronic administration of high-dose corticosterone (CORT) throughout the postpartum in rodents recapitulates key features of PPD, including disrupted maternal care, increased passive coping, HPA axis dysregulation, and compromised hippocampal integrity [11–16], paralleling findings in individuals with PPD [17–21]. The convergence of clinical and preclinical evidence implicates glucocorticoid dysregulation in PPD and underscores the translational utility of this model for investigating the mechanisms underlying PPD and antidepressant efficacy.

Both neuroplasticity and inflammation are associated with MDD and PND. MDD is associated with lower hippocampal and frontal cortical volume [22–24], hippocampal granule neurons[25], and neurogenesis [26] in both males and female individuals, although most of the studies did not analyze sex as a factor. The postpartum period is associated with reduced hippocampal neurogenesis in rodents [16,27,28]. Chronic stress elicits depressive-like endophenotypes in animal models with reduced hippocampal neurogenesis [29–31] and increases specialized extracellular matrix structures such as perineuronal nets (PNNs). PNNs inhibit neuroplasticity [32,33] in cortical and limbic regions [34–36], although again most research has been conducted in males. One study found that PNNs in the medial preoptic area are increased during gestation and decreased in the postpartum period [37], but studies have not examined PNNs in limbic regions associated with PPD. The maternal immune system undergoes significant changes during the perinatal period [38–41] and higher expression of IL-6, IL-1β, and TNF-α in the postpartum is associated with maternal depressive symptoms [42–44]. Neuroplasticity is also related to immune system signalling, with IL-1β reducing neurogenesis in the hippocampus [45–48]. Thus, there is a need to further investigate the role of immune signalling and how it may be linked to neuroplasticity in PPD.

SSRIs remain the first-line pharmacological treatment for PND [49–51] and meta-analyses examining the efficacy of SSRIs show some symptom improvement [51]. However, therapeutic efficacy is heterogeneous [52–54], and most studies do not systematically compare timing of onset or early versus late postpartum response rates. One study found benefits of SSRIs within 4 weeks of treatment, but this benefit diminished at 18-weeks [54]. A similar effect is observed in our rodent model of PPD, as SSRIs improve the maternal care deficits early in the postpartum but do not improve other endophenotypes in the later postpartum (increased passive coping, reductions in hippocampal plasticity) [13,15,16], possibly indicating a lack of efficacy of SSRIs long-term in the postpartum, mirroring human findings [54]. Interestingly, inflammatory signaling is associated with antidepressant efficacy in individuals with MDD, as non-responders to SSRIs showed elevations in peripheral IL-1β levels [55–59]. Previously, we found that the SSRI fluoxetine (FLX) increased hippocampal IL-1β in the postpartum, which coincided with a lack of efficacy in rescuing the CORT-induced increase in passive coping [15]. Notably, increased IL-1β levels activate the serotonin transporter, therefore inducing serotonin re-uptake, which may explain how IL-1β could be mediating reduced SSRI efficacy [60,61].

In the current study, we examined the effects of blocking IL-1β signaling concurrent with FLX treatment in a rodent model of PPD. A high dose of CORT was administered to induce depressive-like endophenotypes in postpartum dams. A subset of dams were additionally given FLX and/or Anakinra (KIN), an IL-1 receptor (IL-1R) antagonist. Outcome measures included depressive-like endophenotypes (maternal care behaviours, passive-coping behaviour, and neuroplasticity (PNNs) in the hippocampus and frontal cortex as well as neurogenesis in the hippocampus. We hypothesized that postpartum CORT treatment would induce depressive-like endophenotypes, including disruptions in maternal care behaviour, increased passive coping behaviour, and reduced neuroplasticity. We also hypothesized that postpartum FLX efficacy on depressive-like endophenotypes would be improved by blocking the IL-1R.

## MATERIALS AND METHODS

### Animals

Fifty-eight adult female and thirty-three male Sprague Dawley rats (3 months old) were purchased from Charles River (Montreal, QC, Canada). Females were randomly assigned to one of four experimental groups (described below), and males were utilized only as breeders. Breeding and culling protocols were followed as previously described [15]. For detailed breeding information, see Method S2 in the supplementary information. All protocols were in accordance with ethical guidelines set by the Canadian Council for Animal Care and were approved by the University of British Columbia Animal Care Committee. For specific animal housing information, see Method S1 in the supplementary information.

**Figure. 1.**
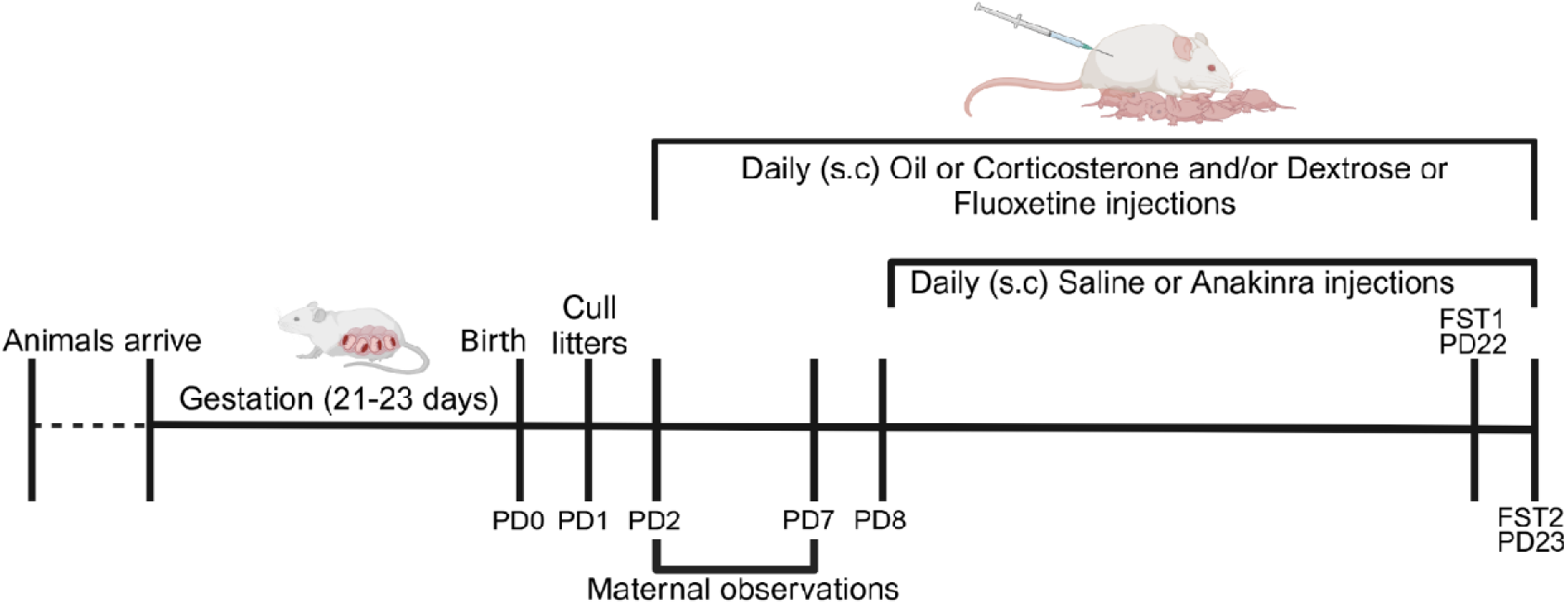
Timeline of experiment. Briefly, animals were bred within the animal facility and kept undisturbed throughout gestation. Date of birth was designated postpartum day (PD) 0. On PD2 animals received daily injections of either corticosterone (CORT) or oil vehicle and fluoxetine (FLX) or dextrose vehicle for 21 days. Maternal care observations occurred during PD2-7. On PD8, animals additionally received daily injections of either anakinra (KIN) or saline (SAL) for 15 days. Forced-swim test (FST) was conducted on PD22 and PD23. Pups remained with the dams until experimental endpoint.

### Drug preparations and treatments

All dams received daily subcutaneous (s.c.) injections beginning from PD2 to PD23 between 8:00–10:00. Dams received corticosterone (CORT; 40 mg/kg, s.c; Sigma-Aldrich, St. Louis, MO, USA) or its vehicle (OIL; sesame oil + 10% EtOH, 1 ml/kg, s.c.) and either fluoxetine (FLX; 10 mg/kg, s.c.; Sequoia Research Products, Pangbourne, UK) or its vehicle, dextrose (DXT; 5% dextrose in sterile water, 1 ml/kg, s.c.). Following the end of the early postpartum, on PD8, dams received an addition of either Anakinra (KIN;100 mg/kg; s.c; Kineret, Amgen, Thousand Oaks, CA, USA) or its vehicle, saline (SAL; 0.9% Sodium Chloride Saline Solution, Baxter, Deerfield, IL, USA). Thus, all dams were randomly assigned to one of four treatment groups (OIL-DXT-S, n=12; CORT-DXT-S, n=13; CORT-FLX-S, n=10; CORT-FLX-KIN, n=11). CORT and FLX were prepared as previously described [11,13,15]. For more information on treatments as well as data from the control groups of OIL-DXT-KIN, OIL-DXT-FLX, OIL-FLX-KIN, CORT-DXT-KIN see Method S3 in the supplementary information.

### Maternal care behavior observations

As previously described [11,13,15], maternal care behavior observations were conducted twice per day (12:00-13:00 and 16:30–17:30) for six days during PD2-7 in the home cage by trained observers. Observations occurred at least 2Lh after injections and after the previous observation. Each observation per animal lasted for 20Lmin per day (10Lmin per morning and afternoon session).The following maternal care behaviors were scored: (a) nursing, which included arched-back nursing (arched-back, actively nursing pups), blanket nursing (in nursing position but resting on-top of pups), and passive nursing (lying on the side and nursing pups); (b) licking (licking pups while not nursing any other pups), (c) self-grooming (not engaging with pups, and self-grooming), (d) nest building (actively building nest), and (e) time spent off-nest (off-nest and not engaging with pups).

### Forced-swim test

As previously described [11,13,15], the forced swim test (FST) was used to assess passive-coping behaviors as a reliable measure of antidepressant efficacy, particularly in rat models [62]. Dams were tested on PD22 (FST1) for 15 minutes, and PD23 (FST2) for 5 minutes. Videos were manually scored using BORIS software (University of Torino, Torino, IT) (Friard and Gamba, 2016) for swimming, climbing, and immobility. For more information on the FST protocol, see Method S4 in the supplementary information.

### Estrous cycle cytology

To examine any influence of estrous stage on behavior, vaginal lavage samples were taken prior to FST2 and were counterstained using cresyl-violet solution and staged based on [63].

### Blood and tissue collection

Immediately following FST2, dams were euthanized by rapid decapitation, and trunk blood was collected. Brains and adrenal glands were removed on a cold surface, and adrenal glands were weighed. Brains were extracted and cut longitudinally into halves. The left hemispheres of each brain were micro dissected into the dorsal hippocampus, ventral hippocampus, and frontal cortex. Dissected tissue was flash-frozen on dry ice and stored at −80 °C for biochemical analyses. Right hemispheres were post-fixed and processed for histological analyses. Full tissue-processing procedures are provided in the Supplement Method S5.

### Immunohistochemistry

All immunohistochemical procedures were conducted on free-floating brain sections for both doublecortin (DCX), a microtubule-associated protein expressed in immature neurons [64], and wisteria floribunda lectin (WFA), a marker of perineuronal nets [32,33]. Sections were permeabilized, blocked, and incubated with primary antibodies for 24 (WFA) and 48 hours (DCX). For WFA staining, fluorescein (FITC) was directly conjugated in the WFA-primary antibody. For the second day of DCX staining, sections were incubated in the secondary antibody for DCX. Tissue used in both stains was counterstained with DAPI, mounted onto Superfrost Plus slides, and cover slipped with anti-fade medium. Complete reagent details, antibody information, and staining protocols are provided in Supplementary Method S6.

### Microscopy and cell quantification

All brain regions were defined according to a standard rat brain atlas (Paxinos and Watson, 2004) and traced using ImageJ software (Image J, 2024). The regions of interest included the dorsal hippocampus (dHPC) and the ventral hippocampus (vHPC). In the frontal cortex, the anterior cingulate cortex (ACC), the prelimbic cortex (PrL), and the infralimbic cortex (IL). Images were acquired on a Zeiss Axio Scan 7 (Carl Zeiss Microscopy, Thornwood, NY, USA) with a 20x objective lens. An investigator blinded to experimental conditions quantified DCX-and WFA-positive (DCX+, WFA+) cells. Density of positive cells was calculated by dividing the total number of cells by area of the corresponding region (cells/mm^2^). DCX+ cells were exhaustively counted manually from 2 sections of dorsal dentate gyrus and 2 sections of ventral dentate gyrus using ZEISS Zen Blue 3.10 Software (Carl Zeiss Microscopy, Thornwood, NY, USA). WFA+ cells were exhaustively counted manually from 2 sections for hippocampal regions (dHPC, vHPC) and medial frontal cortex regions (ACC, PrL, and IL) using ImageJ software (Image J, 2024).

### Tissue homogenization

Following previous protocols (Bodnar et al., 2017; Eid et al., 2019; Qiu et al., 2020), briefly, all tissues were homogenized in 200-400μl cold lysis buffer (150LmM NaCl, 20LmM Tris, pH 7.5, 1LmM EDTA, 1LmM EGTA, 1% Triton X, containing a cocktail of protease and phosphatase inhibitors) using the Qiagen TissueLyserII Lysing Machine (Qiagen, Hilden, Germany). Immediately after homogenization, all tissue samples were centrifuged at 4L°C at 1000xg for 10 min. Supernatant was then collected and stored in −80□°C for protein and analysis.

### PSD-95 and protein quantification

Postsynaptic density-95 (PSD-95) concentrations were measured as previously described [65] using the PSD-95 kit from Meso Scale Discovery (Rockville, MD, USA) according to the manufacturer’s instructions. Samples were run in duplicates to quantify PSD-95 in each sample. Plates were read using a Sector Imager 2400 (Meso Scale Discovery), and data analyses were conducted using the Discovery Workbench 4.0 software (Meso Scale Discovery).Total protein levels were quantified in tissue homogenates using the Pierce Microplate BCA Protein Assay Kit in triplicates following the manufacturer’s protocol (Thermo Scientific Pierce Protein Biology, Thermo Fisher Scientific, Waltham, MA, USA). PSD-95 levels in the HPC and frontal cortex were normalized to total protein levels and reported as PSD-95/mg of protein.

### Statistical analyses

Statistical analyses were performed using Statistica (v. 9, StatSoft, Inc., Tulsa, OK, USA). A one-way factorial analysis of variance (ANOVA) was used on dependent variables of interest (immobility in the FST and PNNs) with treatments (OIL-DXT-SAL, CORT-DXT-SAL, CORT-FLX-SAL, and CORT-FLX-KIN) as the between-subject factor. For nursing behaviour, between-subjects variables were CORT (CORT or OIL) and FLX (FLX or SAL) as KIN administration began after maternal observations ended (PD8). For hippocampal PNNs and DCX+ cells, repeated measures ANOVAs were run across regions (dorsal and ventral dentate gyrus) as within-subjects.Post-hoc tests used Newman-Keuls. *A priori* comparisons were subjected to Bonferroni correction. Effect sizes were reported for significant effects as □_p_^2^□or Cohen’s d.

Effects were considered statistically significant if p□≤□0.05.

## Results

### Postpartum CORT treatment reduced body mass and relative adrenal mass

Postpartum CORT, regardless of adjuvant treatments, reduced weight gain relative to oil-treated dams with a significant decrease in weight from baseline (PD2) starting PD19 (*p*=0.005, Cohen’s d=0.656; Day by Treatments interaction, *F*(21,882)□= 5.736, *p*l<□0.001, □_p_^2^□=0.29). There was also a main effect of DAY, *F*(21,882)□= 17.496, *p*l<□0.001, □_p_^2^□=0.294) but no other significant effects or interactions (all p’s□≥□0.318).

Postpartum CORT treatment significantly reduced relative adrenal mass compared to oil-treated dams (main effect of Treatment; *F*(1,32)□=□22.269, *p*□<□0.001, □_p_^2^□=0.676). There was no significant difference in estrous cycle phase between groups at endpoint (χ2□=15.791, *p*=0.0714; Supplemental Figure 1). Using estrous cycle stage as a covariate did not alter any of the findings (all *p*’s□>0.417).

**Figure 2.**
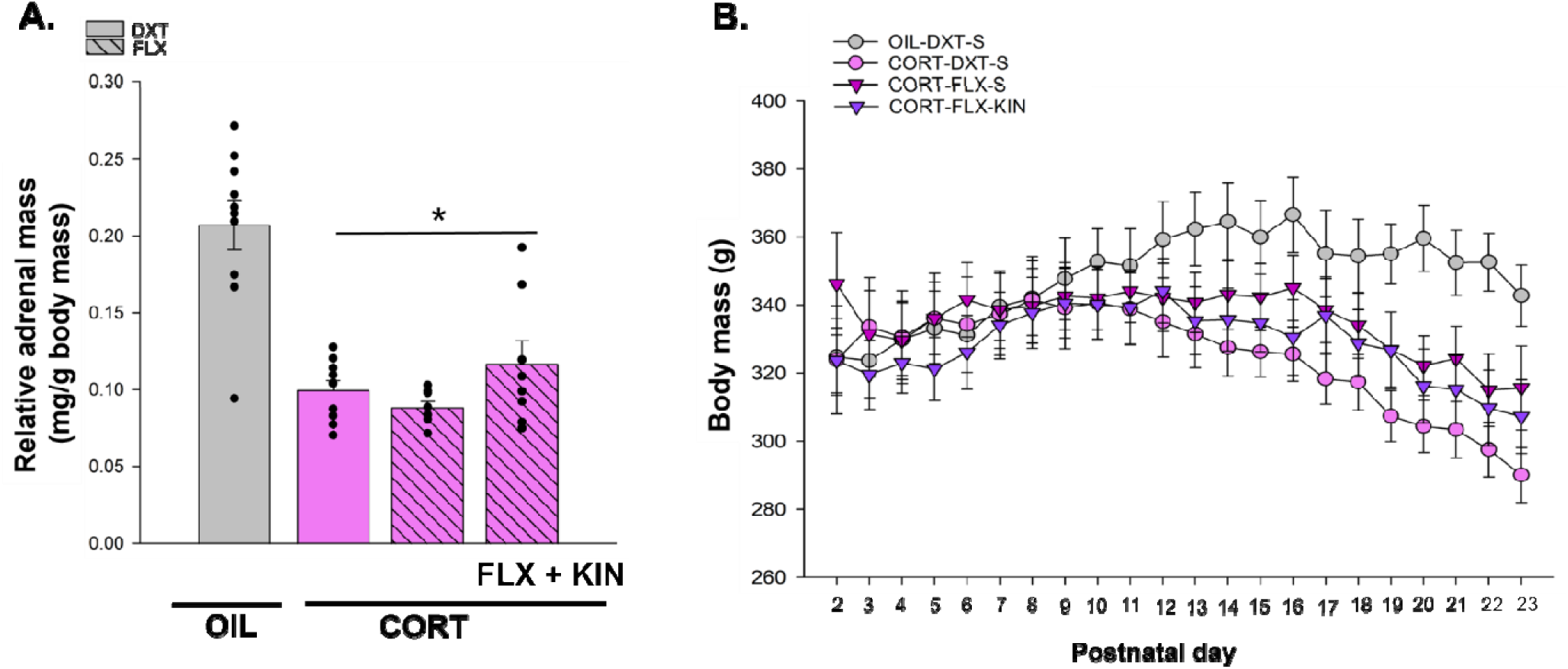
Relative adrenal mass and body mass. **A.** Postpartum CORT treatment significantly reduced relative adrenal mass. *Indicate significance at *p*<0.05. **B.** Postpartum CORT treatment significantly reduced body mass, regardless of adjuvant treatment *Indicate significance at *p*<0.05. n’s = 10-13 per group.

### Postpartum CORT reduced total time nursing and increased total time off nest, Postpartum FLX reversed the effects of Postpartum CORT on maternal behaviours

To investigate the ability of FLX and KIN to rescue PPD-like behavior, several measures of maternal behavior were assessed (Figure. 3). Postpartum CORT significantly reduced total time spent nursing compared to oil-treated dams (*p*=0.001, Cohen’s d=1.533; main effect of CORT; *F*(1,61)□= 11.58, *p*□<□0.001, □_p_^2^= 0.159). Postpartum FLX increased nursing relative to CORT (p=0.007, Cohen’s d=1.036) but not oil-treated controls (*p*=0.327; main effect of FLX, *F*(1,61)□= 4.024, *p*□=0.008, □_p_^2^=0.0492). There were no other significant main or interaction effects for nursing behaviours (all p’s>0.064). Postpartum CORT significantly reduced time spent blanket nursing compared to oil-treated controls (*p*=0.0315, Cohen’s d=0.554; main effect of CORT, *F*(1,61)□= 4.199, *p*□=0.0447, □_p_^2^=0.0644). Postpartum FLX increased time spent arched-back nursing relative to postpartum CORT (*p*=0.036, Cohen’s d=0.856) and was not significantly different than oil-treated controls (p=0.327; main effect of FLX, *F*(1,61)□= 4.017, *p*□=0.0494, □_p_^2^=0.061). Postpartum CORT significantly increased time spent licking relative to OIL controls (*p*=0.0352, Cohen’s d=0.165; main effect of CORT *F*(1,61) = 7.777, *p*=0.007, □_p_^2^= 0.113).

**Figure 3.**
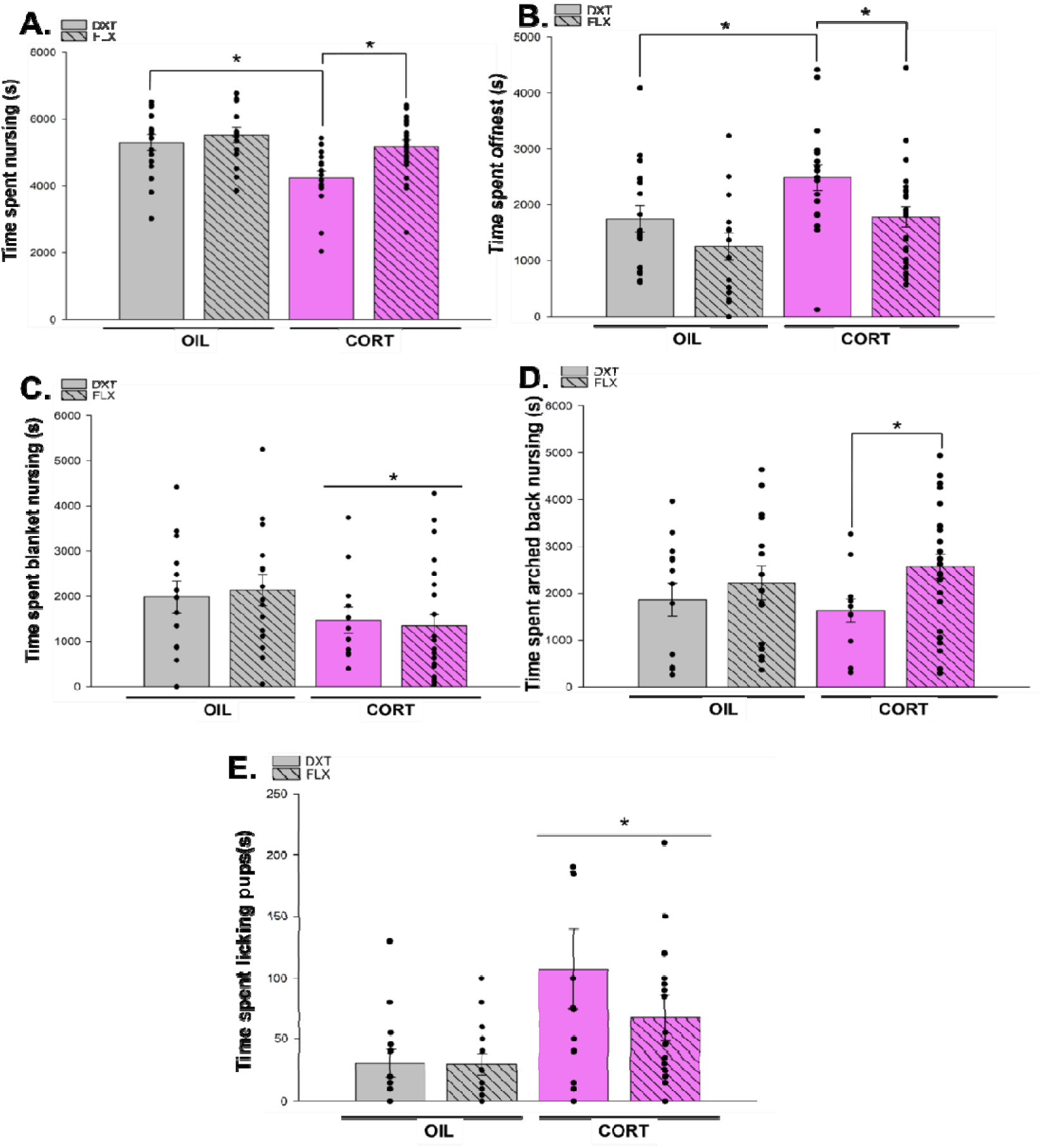
Effects of postpartum corticosterone (CORT) and fluoxetine (FLX) treatment on maternal care behavior (total nursing, total off-nest, blanket nursing, arched back nursing, and licking pups behaviour). **A.** Postpartum CORT treatment significantly reduced total nursing time, and postpartum FLX rescued this reduction. *Indicate significance at *p*<0.05. **B.** Postpartum CORT treatment significantly increased total off-nest time and postpartum FLX rescued this effect. *Indicate significance at *p*<0.05. **C.** Postpartum CORT treatment significantly decreased blanket back nursing *Indicate significance at *p*<0.05. **D.** Postpartum FLX treatment significantly increased time spent arched back nursing *Indicate significance at *p*<0.05. **E.** Postpartum CORT treatment significantly increased time spent licking pups *Indicate significance at *p*<0.05. Data represented in means +standard error of the mean (SEM), overlaid with individual data points. n’s = 10-13 per group.

Postpartum CORT significantly increased total time spent off-nest compared to oil-treated dams (OIL-DXT: *p*=0.035, Cohen’s d=0.942; OIL-FLX: *p*=0.004, Cohen’s d=1.183; main effect of CORT; *F*(1,61)□= 8.575, *p*=0.004, □_p_^2^□=□0.123). Postpartum FLX rescued the CORT-induced increase in time spent off-nest relative to CORT-treated dams (a priori, *p*=0.051, Cohen’s d=0.651, main effect of FLX, *F*(1,61)□= 4.083, *p*□=0.047, □_p_^2^=0.062) and was not significantly different compared to oil-treated controls (OIL-DXT: p=0.523). All other main or interaction effects were not significant (*p*’s > =0.5). No other significant effects on other maternal care behaviours were found (all *p*’s > =0.072)

### Postpartum CORT increased time spent immobile in the forced swim test which was not rescued by any adjuvant treatment

Postpartum CORT significantly increased time spent immobile and reduced the latency to immobility compared to oil-treated dams (time spent immobile: *p*=0.013 Cohen’s d=1.176; latency to immobility: *p*>0.001 Cohen’s d=1.975). Neither concurrent postpartum FLX nor anakinra treatments reversed the effects of postpartum CORT (CORT-FLX-S, time spent immobile: *p*=0.67; latency to immobility: *p*=0.767; CORT-FLX-KIN, time spent immobile: *p*=0.93; latency to immobility: *p*=0.833; time spent immobile: main effect of Treatments, *F*(1,39)□= 4.82, *p*=0.0059, □_p_^2^□=0.27; latency to immobility: main effect of Treatments, *F*(1,39)□=9.56, *p*□<□0.001, □_p_^2^□=0.423). There were no other significant effects found for time spent immobile or latency to immobility (all p’s□≥□0.278).

Postpartum CORT decreased time spent mobile compared to oil-treated dams (*p*=0.010, Cohen’s d=1.166), regardless of FLX and anakinra treatments (CORT-FLX-S, *p*= 0.0055, Cohen’s d=1.302; CORT-FLX-KIN, *p*=0.019, Cohen’s d=1.219; main effect of Treatments, *F*(1,39)□= 5.168, *p*=0.004, □p2□=0.284). There were no other significant effects found for time spent mobile in the FST (all p’s□≥□0.237).

**Figure 4.**
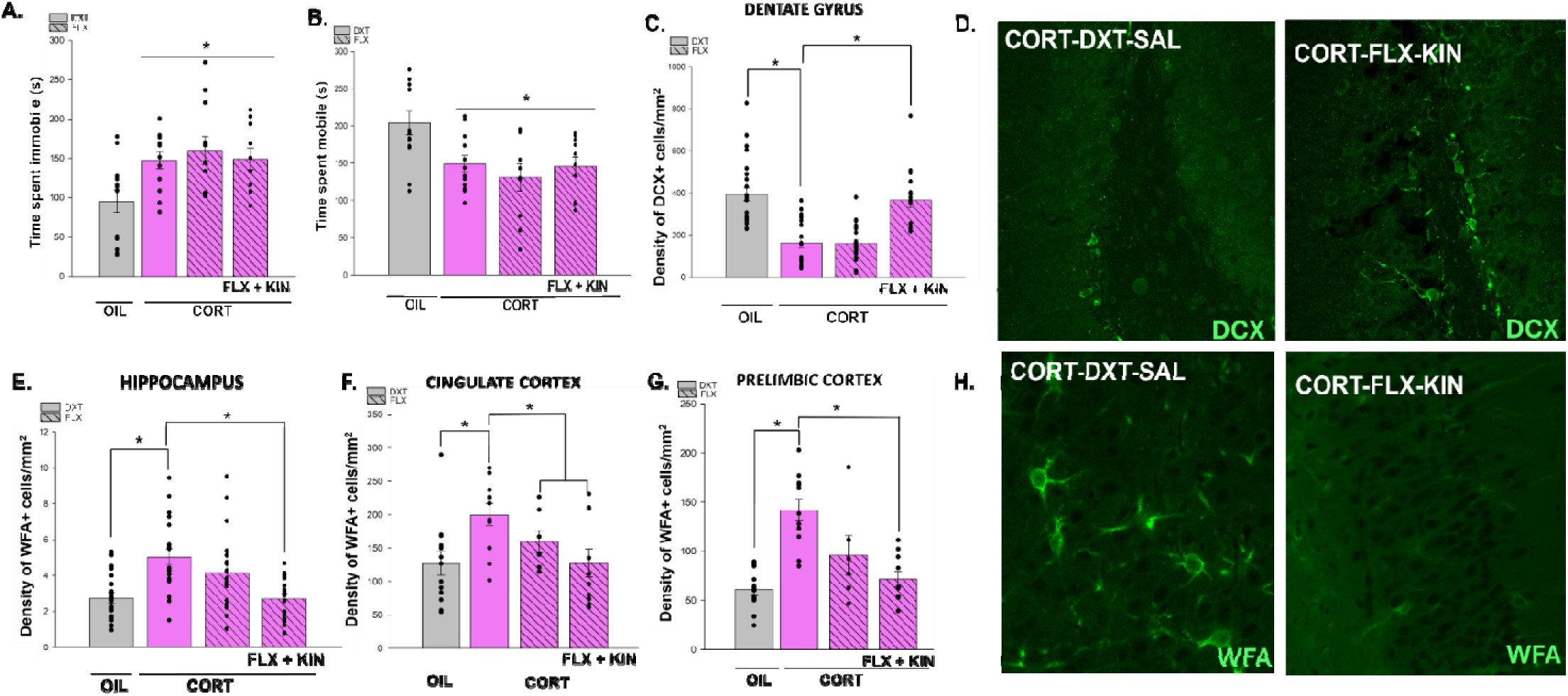
Effects of postpartum CORT, FLX, and anakinra (KIN) on immobility behaviou during the forced swim test (FST), and on doublecortin (DCX)+ cells and wisteria floribunda lectin WFA+ cell density. **A.** Postpartum CORT treatment significantly increased the time spent immobile in the FST. *Indicate significance at *p*<0.05. **B.** Postpartum CORT treatment significantly reduced time spent mobile in the FST. *Indicate significance at *p*<0.05. Data represented in means +standard error of the mean (SEM), overlaid with individual data points. **C.** Postpartum CORT significantly decreased the density of DCX+ cells in the dorsal and ventral hippocampal regions. FLX+KIN rescued these effects. *Indicate significance at *p*<0.05. **D.** Postpartum CORT significantly increased the density of WFA+ cells in the dorsal and ventral hippocampal regions. FLX+KIN rescued these effects. Indicate significance at *p*<0.05. **E.** Postpartum CORT significantly increased the density of WFA+ cells in the cingulate cortex. FLX+KIN rescued these effects. Indicate significance at *p*<0.05. **F.** Postpartum CORT significantly increased the density of WFA+ cells in the prelimbic cortex. Both postpartum FLX+S and FLX+KIN rescued these effects. Indicate significance at *p*<0.05. Data represented in means +standard error of the mean (SEM), overlaid with individual data points. n’s = 10-13 per group. **G.** Representative photomicrographs of CORT-DXT-S, and CORT-FLX-KIN DCX+ cell (green) density taken with 20x objective in the dorsal DG. **H**. Representative photomicrographs of CORT-DXT-S, and CORT-FLX-KIN PNN+ cell (green) density taken with a 20x objective in the CA1 of the hippocampus.

### Postpartum CORT reduced hippocampal neurogenesis, and FLX and KIN administered together rescued the CORT-induced reduction

Postpartum CORT reduced the density of DCX+ cells relative to oil treated dams in both regions of the dentate gyrus (*p*<0.001, Cohen’s d=1.712; Main effect of Treatments *F*(3, 38)□=□36.704, p□<□0.001, □ ^2^□= 0.743). Postpartum FLX also reduced the density of DCX+ cells relative to oil treated dams in both regions of the dentate gyrus (*p*<□0.001, Cohen’s d=1.823). However, CORT-FLX-KIN treatment significantly increased the density of DCX+ cells compared to all other CORT-treated dams in both regions of the dentate gyrus (all *p*’s <0.001; CORT-DXT-S, Cohen’s d=1.621; CORT-FLX-S, Cohen’s d=1.744). There were no significant differences compared to oil-and saline-treated controls (*p*=0.387). The dorsal region had greater density of DCX+ cells than the ventral region regardless of treatment (Main effect of Region *F*(1, 38)□=□70.455, *p*□<□0.001, □ ^2^□= 0.649). There were no other significant interaction effects (all *p*’s>0.457). Supplemental Figure 2 shows all control groups, with separate dorsal and ventral hippocampus figures.

### Postpartum CORT increased the density of WFA+ cells in the hippocampus, which was rescued by FLX and KIN administered together

Postpartum CORT increased the density of WFA+ cells relative to oil-treated dams in both dorsal and ventral hippocampus regions (p<0.001; Cohen’s d=1.33, Main effect of Treatments *F*(3, 34)□=□8.95, *p*□<□0.001, □p2□= 0.441). Postpartum FLX also increased the density of WFA+ cells relative to oil-treated dams in both hippocampal regions (*p*<□0.001, Cohen’s d=0.80). However, CORT-FLX-KIN treatment significantly decreased the density of WFA+ cells compared to CORT-treated dams (all *p*’s <0.001; Cohen’s d=0.832-1.378) and did not differ from oil-treated dams (*p*=0.826). Furthermore, there was a higher density of WFA+ cells in the ventral compared to dorsal hippocampus (main effect of Region, *F*(1, 34□=91.373, *p*<□0.001, □_p_^2^□= 0.728). There was no significant interaction effect (all *p*’s >0.081). Supplemental Figure 4 shows all control groups, with separate dorsal and ventral hippocampus figures.

### Postpartum CORT increased density of WFA+ cells in the ACC and prelimbic cortex, which was rescued by FLX and KIN administered together

Postpartum CORT increased WFA+ cells in the ACC (p=0.004, Cohen’s d=1.229; Main effect of Treatment *F*(3, 88)□= 2.950, *p*□=0.017, □p2=0.11) and PrL(*p*<0.001, Cohen’s d=2.681; Main effect of Treatment *F*(3, 35)□= 13.949, *p*<0.001, □_p_^2^=0.544) relative to oil-treated dams. In the ACC, FLX and KIN together (*p*=0.007, Cohen’s d=1.199) in CORT treated groups significantly decreased WFA+ cell density compared to controls (CORT-DXT-S). In the PrL, both FLX alone (*p*=0.007, Cohen’s d=1.04), and FLX and KIN together (p<0.001, Cohen’s d=2.219) significantly decreased WFA+ cell density compared to controls (CORT-DXT-S). There were no significant differences in the infralimbic cortex (all *p*’s >0.645). Supplemental Figure 4 shows all control groups.

### Postpartum treatments did not influence PSD-95 levels in the hippocampus or frontal cortex

PSD-95 levels were higher in the ventral compared to dorsal hippocampus (main effect of Region in the hippocampus with (*F*(1, 34)□= 8.831, *p*□=0.005, □_p_^2^=0.206). There were no other significant main or interaction effects on PSD-95 levels in either the hippocampus (p>0.096) or frontal cortex (all p’s >0.783).

## Discussion

In the present study, we show that IL-1 receptor antagonism in the postpartum augments the effects of fluoxetine to rescue postpartum CORT-induced reductions in neuroplasticity. Consistent with previous studies, postpartum corticosterone alone increased passive coping, reduced maternal care behaviours, and reduced hippocampal neurogenesis [11–13,15]. Postpartum fluoxetine alone rescued the effects of corticosterone on maternal behaviours, but did not significantly influence any other outcome measures, consistent with past studies [11–13,15]. However, when combined with anakinra, an IL-1R antagonist, fluoxetine treatment rescued the corticosterone-induced changes to neuroplasticity (DCX, WFA) in the postpartum. However, this combination failed to reverse the effects of postpartum corticosterone on passive coping behaviour in the FST, suggesting differential mechanisms drive changes to neuroplasticity versus passive coping. These data enhance our understanding of how manipulation of immune signalling can augment the ability of fluoxetine to increase neuroplasticity in the postpartum, and may be relevant for development of therapeutics for PPD.

### Postpartum corticosterone reduced hippocampal neurogenesis and combined anakinra–fluoxetine treatment restored levels of neurogenesis in the hippocampus

Postpartum corticosterone reduced neurogenesis in the dentate gyrus, consistent with previous work [11,12,16]. Although postpartum fluoxetine was unable to increase neurogenesis alone, anakinra given in conjunction with fluoxetine reversed the postpartum corticosterone reductions in neurogenesis. Moreover, anakinra alone did not rescue the corticosterone-induced reductions in neurogenesis (as shown in Figure S3). These findings point to an important component of SSRI efficacy which may be contingent on the inflammatory milieu [66–68]. SSRIs increase hippocampal neurogenesis [26,69], and, conversely, high levels of IL-1β can reduce hippocampal neurogenesis [45,46]. In male rodent studies, direct IL-1β administration increases depressive-like behaviour [48] and blocking IL-1 receptors also blocks the effects of stress on neurogenesis and anhedonia [70]. Thus, our findings match this existing literature, extending it to postpartum females.

In MDD, non-responders to SSRIs showed elevations in IL-1β levels, whereas SSRI responders do not [55]. Further, increased IL-1β levels activate the serotonin transporter, therefore inducing serotonin re-uptake, which may explain how IL-1β could be reducing the ability of SSRIs to increase neurogenesis [60,61]. Intriguingly, maternal exercise with fluoxetine was also able to reverse the corticosterone-induced reduction in neurogenesis [13], and exercise can reduce IL-1β [71,72], pointing to a consistent role for IL-1β influencing the neurogenic effects of SSRIs. These findings collectively point to why manipulating the IL-1R could increase neurogenesis with fluoxetine.

Studies show that IL-1β influences neurogenesis simultaneously with, and via direct crosstalk with the kynurenine pathway [47]. IL-1β can increase the conversion of tryptophan into its downstream metabolite, kynurenine [73]. Elevated kynurenic acid and other neurotoxic metabolites of the kynurenine pathway have been implicated in PPD in humans [74–76] and in rodents [14,15]. The increase in kynurenine pathway neuro-toxic metabolites also decreases neuroplasticity [77]. Given that both IL-1β and kynurenine metabolites reduce neuroplasticity, both systems may compound to exert these effects in PPD. The unique interplay of immune, and tryptophan metabolism cross-talk emphasizes the need for targeting multiple systems in treating such a heterogeneous disease such as PPD. Future work on the specific mechanisms of these systems are needed within the postpartum context.

### Postpartum corticosterone elevated perineuronal net density and co-administration of anakinra and fluoxetine reversed this effect in the hippocampus, anterior cingulate and prelimbic regions of the frontal cortex

In the present study, postpartum corticosterone increased the density of PNNs in the hippocampus, anterior cingulate, and prelimbic regions, which suggests decreased neuroplasticity throughout these regions. Interestingly, all of these regions are involved in HPA axis homeostasis and MDD, due to glucocorticoid receptors in these regions [30,78]. Glucocorticoids modulate the expression of PNNs by modulating the extracellular matrix components neurocan and hyaluronan synthase [79,80]. Thus, the increase in PNNs may be reducing plasticity in these regions, and limiting glucocorticoid receptors from inhibiting the corticosterone-induced stress response. In the present study there were no significant effects of postpartum corticosterone on the infralimbic cortex to increase PNNs, however, the infralimbic cortex has divergent actions on the stress response compared to the anterior cingulate cortex and prelimbic cortex [81–83].

Chronic stress increases PNN density in the hippocampus and medial prefrontal cortex in male rodents [34–36,84], and chronic fluoxetine treatment can reduce PNN density in the hippocampus, and medial frontal cortex in males [85,86]. In our study, fluoxetine alone was only able to reduce PNNs in the prelimbic cortex, but not in the hippocampus and anterior cingulate cortex. Similarly, anakinra alone did not rescue the corticosterone induced increase in PNNs in the hippocampus, but did rescue these effects in the prelimbic and cingulate cortices (as shown in Figure S4). In the hippocampus, PNNs required both fluoxetine and anakinra to rescue the corticosterone induced increase in PNNs. However, in the cingulate and prelimbic cortices, anakinra alone also sufficed in reducing PNN’s. There may be functional differences in PNNs across regions. PNNs are regulated by glial cells such as astrocytes and microglia [87,88] and microglia are involved in IL-1 signalling [83], thus it is also possible that there are functional regional differences of glial cells which may also contribute to the differences we saw across regions.

Interestingly, anakinra in conjunction with fluoxetine reversed the corticosterone-induced increase in PNNs in the hippocampus, as well as the prelimbic and cingulate cortices. Mirroring the efficacy of anakinra and fluoxetine together on neurogenesis. Recent work shows that the effects of stress on PNNs may be dependent on IL-1 signalling in the hippocampus in males [90]. In the postpartum, neuroplasticity may be dampened by both serotonin and IL-1R signalling, indicating the importance of the co-treatment. Future work should examine the potential mechanisms behind the different effects of treatments across regions. Nevertheless, these findings point to fluoxetine and anakinra together potentially having a more robust effect on PNNs than fluoxetine alone.

### Postpartum corticosterone reduced maternal care and increased passive coping behaviour

Consistent with previous studies, postpartum corticosterone significantly increased time spent passive coping in the FST, and reduced nursing behaviours in the dams. Although fluoxetine rescued nursing behaviours in the early postpartum (PD2-7), it did not reduce passive coping in the late postpartum (PD23), which is also consistent with previous studies [11–13,15]. Notably, maternal care behaviours such as nursing are highly driven by the serotonergic system, indirectly and directly [91–93]. Hence, it is perhaps not surprising that fluoxetine had potent effects on maternal behaviours in the early postpartum.

Our study also showed that postpartum corticosterone increased immobility and reduced latency to first immobility but that postpartum fluoxetine alone or with anakinra did not prevent this increase in immobility. This is inconsistent with previous work in males, showing that while lipopolysaccharide-induced increase in IL-1β resulted in higher immobility in the FST, this effect is not seen in transgenic mice with the IL-1R deletion [61]. Gonadectomy can limit the effects of SSRIs on passive coping and IL-1β on depressive-like endophenotypes [94–96]. Given that the postpartum is marked by a hypogonadal state, this hormonal milieu likely impacts the efficacy of fluoxetine in our model of PPD, adding further complexity to the already present immune, and tryptophan signalling crosstalk.

In humans, individuals with postpartum depression take longer to respond to antidepressants and may require more than one antidepressant medication at time of response [49,52,97], which matches our preclinical model showing the need for adjuvant treatment methods. Additionally, given that our previous work found that exercise alone and with fluoxetine was able to rescue neurogenesis, and passive coping [13], future work should identify mechanisms of exercise in postpartum females.

## Conclusions

Here, we demonstrate that combined treatment with anakinra, an IL-1 receptor antagonist, and fluoxetine, an SSRI, reversed corticosterone-induced deficits in neuroplasticity, whereas neither treatment alone was effective. Notably, the restoration of neuroplasticity occurred in the absence of improvements in passive coping behaviour, highlighting a dissociation between the neuroplastic and behavioural responses to treatment. To our knowledge, this is the first study to examine the combined effects of IL-1 receptor antagonism and SSRI treatment in a preclinical model of PPD. Our findings suggest that IL-1 signalling may constrain antidepressant-induced neuroplasticity in the postpartum brain, identifying a potential mechanism underlying limited antidepressant efficacy. Given the heterogeneity of PPD and the limited efficacy of existing treatments, identifying the biological mechanisms that contribute to individual differences in treatment response is critical. Targeting neuroimmune signalling may offer a promising avenue for developing mechanism-based adjunctive treatments and advancing precision medicine for PPD.

## Author contributions

**Romina Garcia de leon:** Data curation, Formal analysis, Investigation, Methodology, Project administration, Visualization, Writing - original draft, Writing - review & editing. **Wansu Qiu:** Project administration, Investigation, Conceptualization. **Phoenix Au-Yeung:** Data curation, Investigation. **Stephanie E Lieblich:** Project administration, Investigation, Conceptualization. **Amanda Namchuk:** Data curation, Investigation**. Liisa A.M. Galea:** Conceptualization, Formal analysis, Funding acquisition, Project administration, Resources, Supervision, Writing – review & editing.

## Acknowledgements

We gratefully acknowledge Muna Ibrahim, Kimberly Ann Go, and Rebecca Rechlin for their help with animal husbandry, and Catherine Rodriguez, Bhakti Almoula, and Niki Shahraki for their assistance in data collection and preliminary analyses.

We would also like to thank all the wonderful staff members at the Centre for Disease Modelling animal facility at the University of British Columbia for the continuous support and help throughout the project.

## References

1. American Psychiatric Association. Diagnostic and Statistical Manual of Mental Disorders Fifth Edition - DSM-5. 5th ed. American Psychiatric Association Publishing; 2013.

2. Qiu W, Hodges TE, Clark EL, Blankers SA, Galea LAM. Perinatal depression: Heterogeneity of disease and in animal models. Front Neuroendocrinol. 2020;59:100854.

3. O’Hara MW, Wisner KL. Perinatal mental illness: Definition, description and aetiology. Best Pract Res Clin Obstet Gynaecol. 2014;28:3–12.

4. Munk-Olsen T, Laursen TM, Pedersen CB, Mors O, Mortensen PB. New parents and mental disorders: a population-based register study. JAMA. 2006;296:2582–2589.

5. Shorey S, Chee CYI, Ng ED, Chan YH, Tam WWS, Chong YS. Prevalence and incidence of postpartum depression among healthy mothers: A systematic review and meta-analysis. J Psychiatr Res. 2018;104:235–248.

6. Molenaar NM, Maegbaek ML, Rommel A-S, Ibroci E, Liu X, Munk-Olsen T, et al. The incidence of depressive episodes is different before, during, and after pregnancy: A population-based study. J Affect Disord. 2023;322:273–276.

7. Kiewa J, Meltzer-Brody S, Milgrom J, Bennett E, Mackle T, Guintivano J, et al. Lifetime prevalence and correlates of perinatal depression in a case-cohort study of depression. BMJ Open. 2022;12:e059300.

8. Fish-Williamson A, Hahn-Holbrook J. Nutritional factors and cross-national postpartum depression prevalence: an updated meta-analysis and meta-regression of 412 studies from 46 countries. Front Psychiatry. 2023;14:1193490.

9. Wang Z, Liu J, Shuai H, Cai Z, Fu X, Liu Y, et al. Mapping global prevalence of depression among postpartum women. Transl Psychiatry. 2021;11:543.

10. Iliadis SI, Comasco E, Sylvén S, Hellgren C, Poromaa IS, Skalkidou A. Prenatal and Postpartum Evening Salivary Cortisol Levels in Association with Peripartum Depressive Symptoms. PLOS ONE. 2015;10:e0135471.

11. Workman JL, Brummelte S, Galea L a. M. Postpartum corticosterone administration reduces dendritic complexity and increases the density of mushroom spines of hippocampal CA3 arbours in dams. J Neuroendocrinol. 2013;25:119–130.

12. Brummelte S, Galea LAM. Chronic corticosterone during pregnancy and postpartum affects maternal care, cell proliferation and depressive-like behavior in the dam. Horm Behav. 2010;58:769–779.

13. Gobinath AR, Richardson RJ, Chow C, Workman JL, Lieblich SE, Barr AM, et al. Voluntary running influences the efficacy of fluoxetine in a model of postpartum depression. Neuropharmacology. 2018;128:106–118.

14. Qiu W, Go KA, Lamers Y, Galea LAM. Postpartum corticosterone and fluoxetine shift the tryptophan-kynurenine pathway in dams. Psychoneuroendocrinology. 2021;130:105273.

15. Qiu W, Duarte-Guterman P, Eid RS, Go KA, Lamers Y, Galea LAM. Postpartum fluoxetine increased maternal inflammatory signalling and decreased tryptophan metabolism: Clues for efficacy. Neuropharmacology. 2020;175:108174.

16. Workman JL, Gobinath AR, Kitay NF, Chow C, Brummelte S, Galea LAM. Parity modifies the effects of fluoxetine and corticosterone on behavior, stress reactivity, and hippocampal neurogenesis. Neuropharmacology. 2016;105:443–453.

17. Gazal M, Motta LS, Wiener CD, Fernandes JC, Quevedo LÁ, Jansen K, et al. Brain-Derived Neurotrophic Factor in Post-Partum Depressive Mothers. Neurochem Res. 2012;37:583–587.

18. Taylor A, Glover V, Marks M, Kammerer M. Diurnal pattern of cortisol output in postnatal depression. Psychoneuroendocrinology. 2009;34:1184–1188.

19. Moehler E, Brunner R, Wiebel A, Reck C, Resch F. Maternal depressive symptoms in the postnatal period are associated with long-term impairment of mother-child bonding. Arch Womens Ment Health. 2006;9:273–278.

20. Moses-Kolko EL, Banihashemi L, Hipwell AE. Reduced postpartum hippocampal volume is associated with positive mother-infant caregiving behavior. J Affect Disord. 2021;281:297–302.

21. Gutiérrez-Zotes A, Labad J, Martín-Santos R, García-Esteve L, Gelabert E, Jover M, et al. Coping strategies for postpartum depression: a multi-centric study of 1626 women. Arch Womens Ment Health. 2016;19:455–461.

22. McKinnon MC, Yucel K, Nazarov A, MacQueen GM. A meta-analysis examining clinical predictors of hippocampal volume in patients with major depressive disorder. J Psychiatry Neurosci JPN. 2009;34:41–54.

23. Gosnell SN, Velasquez KM, Molfese DL, Molfese PJ, Madan A, Fowler JC, et al. Prefrontal cortex, temporal cortex, and hippocampus volume are affected in suicidal psychiatric patients. Psychiatry Res Neuroimaging. 2016;256:50–56.

24. Du M-Y, Wu Q-Z, Yue Q, Li J, Liao Y, Kuang W-H, et al. Voxelwise meta-analysis of gray matter reduction in major depressive disorder. Prog Neuropsychopharmacol Biol Psychiatry. 2012;36:11–16.

25. Boldrini M, Santiago AN, Hen R, Dwork AJ, Rosoklija GB, Tamir H, et al. Hippocampal granule neuron number and dentate gyrus volume in antidepressant-treated and untreated major depression. Neuropsychopharmacol Off Publ Am Coll Neuropsychopharmacol. 2013;38:1068–1077.

26. Boldrini M, Underwood MD, Hen R, Rosoklija GB, Dwork AJ, John Mann J, et al. Antidepressants increase neural progenitor cells in the human hippocampus. Neuropsychopharmacology. 2009;34:2376–2389.

27. Leuner B, Mirescu C, Noiman L, Gould E. Maternal experience inhibits the production of immature neurons in the hippocampus during the postpartum period through elevations in adrenal steroids. Hippocampus. 2007;17:434–442.

28. Eid RS, Chaiton JA, Lieblich SE, Bodnar TS, Weinberg J, Galea LAM. Early and late effects of maternal experience on hippocampal neurogenesis, microglia, and the circulating cytokine milieu. Neurobiol Aging. 2019;78:1–17.

29. Radley JJ, Sisti HM, Hao J, Rocher AB, McCall T, Hof PR, et al. Chronic behavioral stress induces apical dendritic reorganization in pyramidal neurons of the medial prefrontal cortex. Neuroscience. 2004;125:1–6.

30. McEwen BS, Nasca C, Gray JD. Stress Effects on Neuronal Structure: Hippocampus, Amygdala, and Prefrontal Cortex. Neuropsychopharmacology. 2016;41:3–23.

31. Cook SC, Wellman CL. Chronic stress alters dendritic morphology in rat medial prefrontal cortex. J Neurobiol. 2004;60:236–248.

32. Carceller H, Gramuntell Y, Klimczak P, Nacher J. Perineuronal Nets: Subtle Structures with Large Implications. The Neuroscientist. 2023;29:569–590.

33. Fawcett JW, Oohashi T, Pizzorusso T. The roles of perineuronal nets and the perinodal extracellular matrix in neuronal function. Nat Rev Neurosci. 2019;20:451–465.

34. Martinez CA, Pantazopoulos H, Gisabella B, Stephens ET, Garteiser J, Del Arco A. Choice impulsivity after repeated social stress is associated with increased perineuronal nets in the medial prefrontal cortex. Sci Rep. 2024;14:7093.

35. Simard S, Coppola G, Rudyk CA, Hayley S, McQuaid RJ, Salmaso N. Profiling changes in cortical astroglial cells following chronic stress. Neuropsychopharmacology. 2018;43:1961–1971.

36. Pesarico AP, Bueno-Fernandez C, Guirado R, Gómez-Climent MÁ, Curto Y, Carceller H, et al. Chronic Stress Modulates Interneuronal Plasticity: Effects on PSA-NCAM and Perineuronal Nets in Cortical and Extracortical Regions. Front Cell Neurosci. 2019;13.

37. Uriarte N, Ferreño M, Méndez D, Nogueira J. Reorganization of perineuronal nets in the medial Preoptic Area during the reproductive cycle in female rats. Sci Rep. 2020;10:5479.

38. Garcia de leon R, Hodges TE, Brown HK, Bodnar TS, Galea LAM. Inflammatory signalling during the perinatal period: Implications for short-and long-term disease risk. Psychoneuroendocrinology. 2025;172:107245.

39. Maes M, Lin A, Ombelet W, Stevens K, Kenis G, De Jongh R, et al. Immune activation in the early puerperium is related to postpartum anxiety and depressive symptoms. Psychoneuroendocrinology. 2000;25:121–137.

40. Bränn E, Edvinsson Å, Rostedt Punga A, Sundström-Poromaa I, Skalkidou A. Inflammatory and anti-inflammatory markers in plasma: from late pregnancy to early postpartum. Sci Rep. 2019;9:1863.

41. Haim A, Julian D, Albin-Brooks C, Brothers HM, Lenz KM, Leuner B. A survey of neuroimmune changes in pregnant and postpartum female rats. Brain Behav Immun. 2017;59:67–78.

42. Corwin EJ, Pajer K, Paul S, Lowe N, Weber M, McCarthy DO. Bidirectional psychoneuroimmune interactions in the early postpartum period influence risk of postpartum depression. Brain Behav Immun. 2015;49:86–93.

43. Silva-Fernandes A, Conde A, Marques M, Caparros-Gonzalez RA, Fransson E, Mesquita AR, et al. Inflammatory biomarkers and perinatal depression: A systematic review. PloS One. 2024;19:e0280612.

44. Boufidou F, Lambrinoudaki I, Argeitis J, Zervas IM, Pliatsika P, Leonardou AA, et al. CSF and plasma cytokines at delivery and postpartum mood disturbances. J Affect Disord. 2009;115:287–292.

45. Wu MD, Hein AM, Moravan MJ, Shaftel SS, Olschowka JA, O’Banion MK. Adult murine hippocampal neurogenesis is inhibited by sustained IL-1β and not rescued by voluntary running. Brain Behav Immun. 2012;26:292–300.

46. O’Léime CS, Cryan JF, Nolan YM. Nuclear deterrents: Intrinsic regulators of IL-1β-induced effects on hippocampal neurogenesis. Brain Behav Immun. 2017;66:394–412.

47. Zunszain PA, Anacker C, Cattaneo A, Choudhury S, Musaelyan K, Myint AM, et al. Interleukin-1β: A New Regulator of the Kynurenine Pathway Affecting Human Hippocampal Neurogenesis. Neuropsychopharmacology. 2012;37:939–949.

48. Goshen I, Kreisel T, Ben-Menachem-Zidon O, Licht T, Weidenfeld J, Ben-Hur T, et al. Brain interleukin-1 mediates chronic stress-induced depression in mice via adrenocortical activation and hippocampal neurogenesis suppression. Mol Psychiatry. 2008;13:717–728.

49. Stewart DE, Vigod SN. Postpartum Depression: Pathophysiology, Treatment, and Emerging Therapeutics. Annu Rev Med. 2019;70:183–196.

50. Cooper WO, Willy ME, Pont SJ, Ray WA. Increasing use of antidepressants in pregnancy. Am J Obstet Gynecol. 2007;196:544.e1–5.

51. Molenaar NM, Kamperman AM, Boyce P, Bergink V. Guidelines on treatment of perinatal depression with antidepressants: An international review. Aust N Z J Psychiatry. 2018;52:320–327.

52. De Crescenzo F, Perelli F, Armando M, Vicari S. Selective serotonin reuptake inhibitors (SSRIs) for post-partum depression (PPD): A systematic review of randomized clinical trials. J Affect Disord. 2014;152–154:39–44.

53. Cox EQ, Sowa NA, Meltzer-Brody SE, Gaynes BN. The Perinatal Depression Treatment Cascade: Baby Steps Toward Improving Outcomes. J Clin Psychiatry. 2016;77:1189–1200.

54. Sharp DJ, Chew-Graham CA, Tylee A, Lewis G, Howard L, Anderson I, et al. A pragmatic randomised controlled trial to compare antidepressants with a community-based psychosocial intervention for the treatment of women with postnatal depression: the RESPOND trial. Health Technol Assess. 2010;14:1–181.

55. Syed SA, Beurel E, Loewenstein DA, Lowell JA, Craighead WE, Dunlop BW, et al. Defective Inflammatory Pathways in Never-Treated Depressed Patients Are Associated with Poor Treatment Response. Neuron. 2018;99:914–924.e3.

56. Hannestad J, DellaGioia N, Bloch M. The Effect of Antidepressant Medication Treatment on Serum Levels of Inflammatory Cytokines: A Meta-Analysis. Neuropsychopharmacology. 2011;36:2452–2459.

57. Uint L, Bastos GM, Thurow HS, Borges JB, Hirata TDC, França JID, et al. Increased levels of plasma IL-1b and BDNF can predict resistant depression patients. Rev Assoc Medica Bras 1992. 2019;65:361–369.

58. Haroon E, Daguanno AW, Woolwine BJ, Goldsmith DR, Baer WM, Wommack EC, et al. Antidepressant treatment resistance is associated with increased inflammatory markers in patients with major depressive disorder. Psychoneuroendocrinology. 2018;95:43–49.

59. Zou W, Feng R, Yang Y. Changes in the serum levels of inflammatory cytokines in antidepressant drug-naïve patients with major depression. PLOS ONE. 2018;13:e0197267.

60. Zhu C-B, Blakely RD, Hewlett WA. The Proinflammatory Cytokines Interleukin-1beta and Tumor Necrosis Factor-Alpha Activate Serotonin Transporters. Neuropsychopharmacology. 2006;31:2121–2131.

61. Zhu C-B, Lindler KM, Owens AW, Daws LC, Blakely RD, Hewlett WA. Interleukin-1 Receptor Activation by Systemic Lipopolysaccharide Induces Behavioral Despair Linked to MAPK Regulation of CNS Serotonin Transporters. Neuropsychopharmacology. 2010;35:2510–2520.

62. Lucki I. The forced swimming test as a model for core and component behavioral effects of antidepressant drugs. Behav Pharmacol. 1997;8:523–532.

63. Cora MC, Kooistra L, Travlos G. Vaginal Cytology of the Laboratory Rat and Mouse: Review and Criteria for the Staging of the Estrous Cycle Using Stained Vaginal Smears. Toxicol Pathol. 2015;43:776–793.

64. Brown JP, Couillard-Després S, Cooper-Kuhn CM, Winkler J, Aigner L, Kuhn HG. Transient expression of doublecortin during adult neurogenesis. J Comp Neurol. 2003;467:1–10.

65. Duarte-Guterman P, Richard JE, Lieblich SE, Eid RS, Lamers Y, Galea LAM. Cellular and molecular signatures of motherhood in the adult and ageing rat brain. Open Biol. 2023;13:230217.

66. Miller AH, Raison CL. The role of inflammation in depression: from evolutionary imperative to modern treatment target. Nat Rev Immunol. 2016;16:22–34.

67. Branchi I, Poggini S, Capuron L, Benedetti F, Poletti S, Tamouza R, et al. Brain-immune crosstalk in the treatment of major depressive disorder. Eur Neuropsychopharmacol. 2021;45:89–107.

68. Nava RG, Adri AS, Filgueiras IS, Nóbile AL, Barcelos PM, Corrêa YLG, et al. Modulation of neuroimmune cytokine networks by antidepressants: implications in mood regulation. Transl Psychiatry. 2025;15:314.

69. Hodes GE, Hill-Smith TE, Suckow RF, Cooper TB, Lucki I. Sex-Specific Effects of Chronic Fluoxetine Treatment on Neuroplasticity and Pharmacokinetics in Mice. J Pharmacol Exp Ther. 2010;332:266–273.

70. Koo JW, Duman RS. Evidence for IL-1 receptor blockade as a therapeutic strategy for the treatment of depression. Curr Opin Investig Drugs Lond Engl 2000. 2009;10:664–671.

71. Ding Y, Xu X. Effects of regular exercise on inflammasome activation-related inflammatory cytokine levels in older adults: a systematic review and meta-analysis. J Sports Sci. 2021;39:2338–2352.

72. Ding Y, Xu X. Anti-inflammatory effect of exercise training through reducing inflammasome activation-related inflammatory cytokine levels in overweight/obese populations: A systematic review and meta-analysis. Complement Ther Clin Pract. 2022;49:101656.

73. Maes M, Leonard BE, Myint AM, Kubera M, Verkerk R. The new ‘5-HT’ hypothesis of depression: Cell-mediated immune activation induces indoleamine 2,3-dioxygenase, which leads to lower plasma tryptophan and an increased synthesis of detrimental tryptophan catabolites (TRYCATs), both of which contribute to the onset of depression. Prog Neuropsychopharmacol Biol Psychiatry. 2011;35:702–721.

74. Duan K-M, Ma J-H, Wang S-Y, Huang Z, Zhou Y, Yu H. The role of tryptophan metabolism in postpartum depression. Metab Brain Dis. 2018;33:647–660.

75. Teshigawara T, Mouri A, Kubo H, Nakamura Y, Shiino T, Okada T, et al. Changes in tryptophan metabolism during pregnancy and postpartum periods: Potential involvement in postpartum depressive symptoms. J Affect Disord. 2019;255:168–176.

76. Achtyes E, Keaton SA, Smart L, Burmeister AR, Heilman PL, Krzyzanowski S, et al. Inflammation and kynurenine pathway dysregulation in post-partum women with severe and suicidal depression. Brain Behav Immun. 2020;83:239–247.

77. de Bartolomeis A, Fornaro M, Scopetta E, Ricci C, Irano A, De Simone G, et al. Chronic stress and brain kynurenine pathway: addressing unresolved issues with a meta-analytic approach of preclinical studies, translational implication for psychiatric disorders. Eur Neuropsychopharmacol. 2025;101:22–40.

78. Diorio D, Viau V, Meaney MJ. The role of the medial prefrontal cortex (cingulate gyrus) in the regulation of hypothalamic-pituitary-adrenal responses to stress. J Neurosci Off J Soc Neurosci. 1993;13:3839–3847.

79. Yue L, Craig MT, Morris BJ. Glucocorticoids Modulate Expression of Perineuronal Net Component Genes and Parvalbumin During Development of Mouse Cortical Neurons. Mol Neurobiol. 2025;63:205.

80. Nguyen HD, Miyazaki H, Kawai H, Wang Z, Sakamoto S, Takaki M, et al. Alteration of perineuronal nets and parvalbumin interneurons in prefrontal cortex and hippocampus, and correlation with blood corticosterone in activity-based anorexia model mice. Neurosci Res. 2025;218:104922.

81. Tao Y, Cai C-Y, Xian J-Y, Kou X-L, Lin Y-H, Qin C, et al. Projections from Infralimbic Cortex to Paraventricular Thalamus Mediate Fear Extinction Retrieval. Neurosci Bull. 2021;37:229–241.

82. Jones KR, Myers B, Herman JP. Stimulation of the Prelimbic Cortex Differentially Modulates Neuroendocrine Responses to Psychogenic and Systemic Stressors. Physiol Behav. 2011;104:266–271.

83. Vidal-Gonzalez I, Vidal-Gonzalez B, Rauch SL, Quirk GJ. Microstimulation reveals opposing influences of prelimbic and infralimbic cortex on the expression of conditioned fear. Learn Mem. 2006;13:728–733.

84. Riga D, Kramvis I, Koskinen MK, van Bokhoven P, van der Harst JE, Heistek TS, et al. Hippocampal extracellular matrix alterations contribute to cognitive impairment associated with a chronic depressive-like state in rats. Sci Transl Med. 2017;9:eaai8753.

85. Ohira K, Takeuchi R, Iwanaga T, Miyakawa T. Chronic fluoxetine treatment reduces parvalbumin expression and perineuronal netsin gamma-aminobutyric acidergic interneurons of the frontal cortex in adultmice. Mol Brain. 2013;6:43.

86. Ueno H, Kitano E, Mori S, Takahashi Y, Murakami S, Wani K, et al. Chronic fluoxetine modulates perineuronal nets and inhibitory neuronal function in relation to behavioral outcomes. IBRO Neurosci Rep. 2025;19:781–791.

87. Wiese S, Karus M, Faissner A. Astrocytes as a Source for Extracellular Matrix Molecules and Cytokines. Front Pharmacol. 2012;3.

88. Crapser JD, Arreola MA, Tsourmas KI, Green KN. Microglia as hackers of the matrix: sculpting synapses and the extracellular space. Cell Mol Immunol. 2021;18:2472–2488.

89. Zhu L, Liu X, Nemeth DP, DiSabato DJ, Witcher KG, Mckim DB, et al. Interleukin-1 causes CNS inflammatory cytokine expression via endothelia-microglia bi-cellular signaling. Brain Behav Immun. 2019;81:292–304.

90. Goodman EJ, Britt RM, Zhao F, Davis AC, Wangler LM, Emami E, et al. Stress Sensitization of Neurons in the Dentate Gyrus Is Dependent on Neuronal Interleukin-1 Receptor Signaling and Is Associated with Increased Synaptic Plasticity, Perineuronal Nets, and Excitatory/Inhibitory Input Imbalance. J Neurosci. 2025;45.

91. Bagdy G. Role of the hypothalamic paraventricular nucleus in 5-HT1A, 5-HT2A and 5-HT2C receptor-mediated oxytocin, prolactin and ACTH/corticosterone responses. Behav Brain Res. 1995;73:277–280.

92. Unroe KA, Maltman JL, Shupe EA, Clinton SM. Disrupted serotonin system development via early life antidepressant exposure impairs maternal care and increases serotonin receptor expression in adult female offspring. Dev Psychobiol. 2022;64:e22292.

93. Angoa-Pérez M, Kane MJ, Sykes CE, Perrine SA, Church MW, Kuhn DM. Brain serotonin determines maternal behavior and offspring survival. Genes Brain Behav. 2014;13:579–591.

94. Gómez ML, Martínez-Mota L, Estrada-Camarena E, Fernández-Guasti A. Influence of the brain sexual differentiation process on despair and antidepressant-like effect of fluoxetine in the rat forced swim test. Neuroscience. 2014;261:11–22.

95. Eid RS, Lieblich SE, Wong SJ, Galea LAM. Ovarian status dictates the neuroinflammatory and behavioral consequences of sub-chronic stress exposure in middle-aged female mice. Neurobiol Stress. 2020;12:100199.

96. Wainwright SR, Lieblich SE, Galea LAM. Hypogonadism predisposes males to the development of behavioural and neuroplastic depressive phenotypes. Psychoneuroendocrinology. 2011;36:1327–1341.

97. Hendrick V, Altshuler L, Strouse T, Grosser S. Postpartum and nonpostpartum depression: differences in presentation and response to pharmacologic treatment. Depress Anxiety. 2000;11:66–72.

98. Latendresse G, Elmore C, Deneris A. Selective Serotonin Reuptake Inhibitors as First-Line Antidepressant Therapy for Perinatal Depression. J Midwifery Womens Health. 2017;62:317–328.

99. Anderson GD, Peterson TC, Vonder Haar C, Kantor ED, Farin FM, Bammler TK, et al. Comparison of the effects of erythropoietin and anakinra on functional recovery and gene expression in a traumatic brain injury model. Front Pharmacol. 2013;4:129.

100. Greenhalgh A, Galea J, Dénes A, Tyrrell P, Rothwell N. Rapid brain penetration of interleukin-1 receptor antagonist in rat cerebral ischaemia: pharmacokinetics, distribution, protection. Br J Pharmacol. 2010;160:153–159.

101. Plümpe T, Ehninger D, Steiner B, Klempin F, Jessberger S, Brandt M, et al. Variability of doublecortin-associated dendrite maturation in adult hippocampal neurogenesis is independent of the regulation of precursor cell proliferation. BMC Neurosci. 2006;7:77.

