## Supplement Methods for "Targeting IL-1 Signaling to Improve Antidepressant Efficacy in a Rodent Model of Postpartum Depression"

**Methods S1.**

**Animals**

All animals were initially housed in same-sex pairs upon arrival in polycarbonate cages (24 × 16 × 46 cm) with aspen chip bedding. Throughout the experiment, rats were maintained on a 12 h: 12 h light/dark cycle (lights on at 07:00), in standard temperature conditions (21 ± 1ºC; 50 ± 10% humidity) and given standard rat chow (Jamieson's Pet Food Distributors Ltd, Delta, BC, Canada) as well as autoclaved tap water *ad libitum*. Males and females were housed in separate colony rooms unless they were breeding. All animals acclimatized to the facilities for seven days and handled for five days prior to the start of the experiment. All protocols were in accordance with ethical guidelines set by the Canadian Council for Animal Care and were approved by the University of British Columbia Animal Care Committee.

**Methods S2.**

**Breeding**

Prior to breeding, males were single-housed for 24h. During breeding, two females and one male were paired at approximately 17:00h and left overnight. The following morning between 8:00-10:00, females were vaginally lavaged to assess for the presence of sperm. Females were deemed pregnant if sperm was detected, and this was considered gestational date (GD)1. Dams were single-housed and placed in a separate colony room and left undisturbed for the duration of gestation, except for weekly cage changes and weighing. Day of birth is regarded as postpartum day (PD) 0. All litters were culled to 5 females and 5 males on PD1 (the following day after birth of the pups and prior to any treatments). A total of 7 dams received cross-fostered pups from aged-matched groups. The first week of postpartum, PD1-7 we define as “early postpartum” and the last week postpartum PD17-PD23, “late postpartum”.

**Methods S3.**

**Drug preparations and treatments**

CORT injections are used over maternal stress models to avoid impacting offspring directly, and model the high basal cortisol seen in PPD [10,18].The CORT dose reliably induces a depressive-like phenotype in dams to reduce maternal behavior, hippocampal neuroplasticity, and increase passive coping behaviours [11–13,15]. The FLX dose alone increases brain derived neurotrophic factor and hippocampal plasticity (Hodes et al., 2010) and improves maternal behavior in conjunction with maternal CORT [11–13,15] and is considered a higher than clinically relevant dose [97]. The KIN (IL-1R antagonist) dose was chosen due to previous work demonstrating therapeutic penetrance of the blood brain barrier in male rats [98,99].

**Methods S4.**

**Forced swim test**

All behavioral testing occurred at least 2h after injections in a separate room from housing. Dams were placed into water tank (11.5 x 25.5 inches) filled to a depth of 17.7 inches containing warm, tap water (24–26 °C) in dim light. Both FST sessions were videotaped and scored by a blinded scorer.

**Methods S5.**

**Blood and tissue collection**

Immediately following FST2, dams were euthanized by rapid decapitation, and trunk blood was collected. Brains and adrenal glands were extracted on a cold surface. Brains were extracted and cut longitudinally into halves. The left hemispheres were microdissected into the dorsal and ventral hippocampus, and frontal cortex. Following dissections, brain tissue was flash frozen on dry ice, and stored at -80ºC. The right hemispheres were postfixed in 4% paraformaldehyde solution for 24h, and then switched to 30% sucrose in 0.1M phosphate buffer saline solution (PBS, pH = 7.4) at 4°C until brains sank. The right hemispheres were post-fixed for 24 h in 4% paraformaldehyde (at 4°C), then transferred to a 30% sucrose solution for cryoprotection until the brains sank. Using a freezing microtome (2M2000R; Leica, Richmond Hill, ON), the right hemisphere of each brain was sliced into 30 μm coronal sections and collected in series of 6 throughout the frontal cortex and in series of 10 throughout the entire rostral-caudal extent of the hippocampus. Sections were stored in a cryoprotective medium (consisting of 0.1 M PBS, 30% ethylene glycol (Sigma-Aldrich), and 20% glycerol (Sigma-Aldrich)) at − 20°C. Adrenal glands were collected and weighed.

**Methods S6.**

**Doublecortin**

Tissue was rinsed in 0.1 M PBS (pH 7.4; 3 × 10 minutes) between each of the following procedures. Tissue was first treated with 10% Triton-X in 0.1M PBS (Sigma Aldrich, St. Louis, MO, USA) for 30 in, then blocked in 10% Normal Donkey Serum (NDS), in 0.3% Triton-X in 0.1M PBS for 1h. Next, tissue was incubated in a primary antibody solution consisting of 1:1000 rabbit anti-DCX (Cell Signaling Technology, Danvers, MA, USA, 4606S) in blocking solution (10% NDS in 0.3% Triton-X in 0.1M PBS) for 48h. Two days later, tissue was incubated in a secondary antibody solution consisting of 1:500 Alexa Fluor 488 donkey anti-rabbit (ThermoFisher, A21206) in 0.1M PBS for 4h. Following, sections were counterstained with DAPI for 3min (14.3 mM solution diluted 1:10000 in 0.1 M PBS, ThermoFisher). Finally, sections were mounted onto Superfrost Plus slides (Fisher Scientific, Inc., Hampton, NH, USA), and cover-slipped with anti-fade medium (0.5 % polyvinyl alcohol-DABCO, Sigma, St. Louis, MO, USA).

**Perineuronal nets**

Tissue was rinsed in 0.1 M PBS (pH 7.4; 3 × 10 minutes) between each of the following procedures. Tissue was first treated with 10% Triton-X in 0.1M PBS (Sigma Aldrich, St. Louis, MO, USA) for 30 in, then blocked in 10% Normal Donkey Serum (NDS), in 0.3% Triton-X in 0.1M PBS for 1h. Next, tissue was incubated in 1:1000 with the plant lectin, conjugated Wisteria Floribunda, fluorescein (FITC) (WFA) antibody solution (ThermoFischer, L32481) in blocking solution (10% NDS in 0.3% Triton-X in 0.1M PBS) for 24h. The following day, sections were counterstained with DAPI for 3min (14.3 mM solution diluted 1:10000 in 0.1 M PBS, ThermoFisher). Finally, sections were mounted onto Superfrost Plus slides (Fisher Scientific, Inc., Hampton, NH, USA), and cover-slipped with anti-fade medium (0.5 % polyvinyl alcohol-DABCO, Sigma, St. Louis, MO, USA).

**Methods S7.**

**Cell Quantification**

To evaluate the rate at which newly produced hippocampal neurons mature, we classified the morphology of DCX+ cells for maturity stage [100]. A total of 50 cells were randomly selected from the dorsal and the ventral hippocampus (2 sections for each). Cells were selected if they were relatively isolated (i.e., not clumped or overlapping) and if they were within the granule cell layer. Cells were classified in one of three developmental stages based on morphological attributes: (1) proliferative if they had no processes or short, plump processes; (2) intermediate if they had medium length processes with no branching; or (3) postmitotic if they had branching dendrites reaching the molecular layer and/or granule cell layer.

**Results S1.**

**Supplemental Table 1. Number of rats in each estrous stage on FST2.**

| **Estrous cycle stage** | **n** | **Percentage** |
|---|---|---|
| Proestrus | 11 | 24.4% |
| Estrus | 10 | 22.2% |
| Metestrus | 15 | 33.3% |
| Diestrus | 9 | 20 |

**Results S2.**

**Postpartum FLX reduced the percentage of postmitotic DCX+ cells in the dorsal dentate gyrus. Postpartum FLX and KIN administered together reduced the percentage of postmitotic DCX+ cells in the ventral dentate gyrus.**

In the dorsal dentate gyrus, postpartum FLX treatment, regardless of KIN, reduced the percentage of postmitotic DCX+ cells compared to all other groups (all *p*'s< 0.034 Cohen's d=0.908-1.634, Stage by Treatment interaction F(6, 92) = 3.519, *p*=0.003, $\eta_p^2$ = 0.186).There were no significant differences among groups in the proliferative or intermediate DCX+ stage type (all *p'*s<0.302). There was also a main effect of Stage *F*(2, 92) = 61.784, *p* < 0.001, $\eta_p^2$= 0.573), but not main effect of treatment (*p*=0.104). In the ventral dentate gyrus, postpartum FLX+KIN treatment reduced the percentage of postmitotic DCX+ cells relative to oil-treated dams (p=0.018, Cohen's d=1.15, Main effect of Treatment, *F*(3, 46) =2.917, *p* =0.044, ηp2= 0.159). There was also a main effect of Stage (Main effect of Stage *F*(2, 92) =24.876, *p* <0.001, $\eta_p^2$= 0.350). There was no significant interaction (all *p*'s ≥ 0.155).

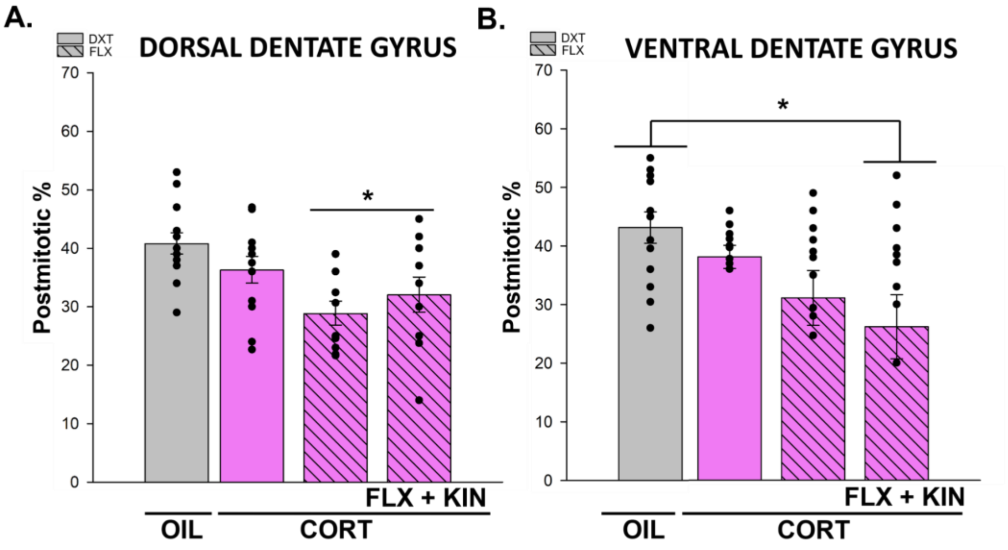


**Supplemental Figure 1. Effects of CORT, FLX, and FLX+ KIN on DCX+ cell density, and DCX+ cells at particular developmental stages. A.** Postpartum FLX significantly decreased the percentage of postmitotic DCX+ cells in the dorsal dentate gyrus. *Indicate significance at $p<0.05$. **B.** Postpartum FLX+KIN treatment significantly decreased the percentage of postmitotic DCX+ cells in the ventral dentate gyrus. *Indicate significance at $p<0.05$. Data represented in means +standard error of the mean (SEM), overlaid with individual data points. n's = 10-13 per group.

**Results S3.**

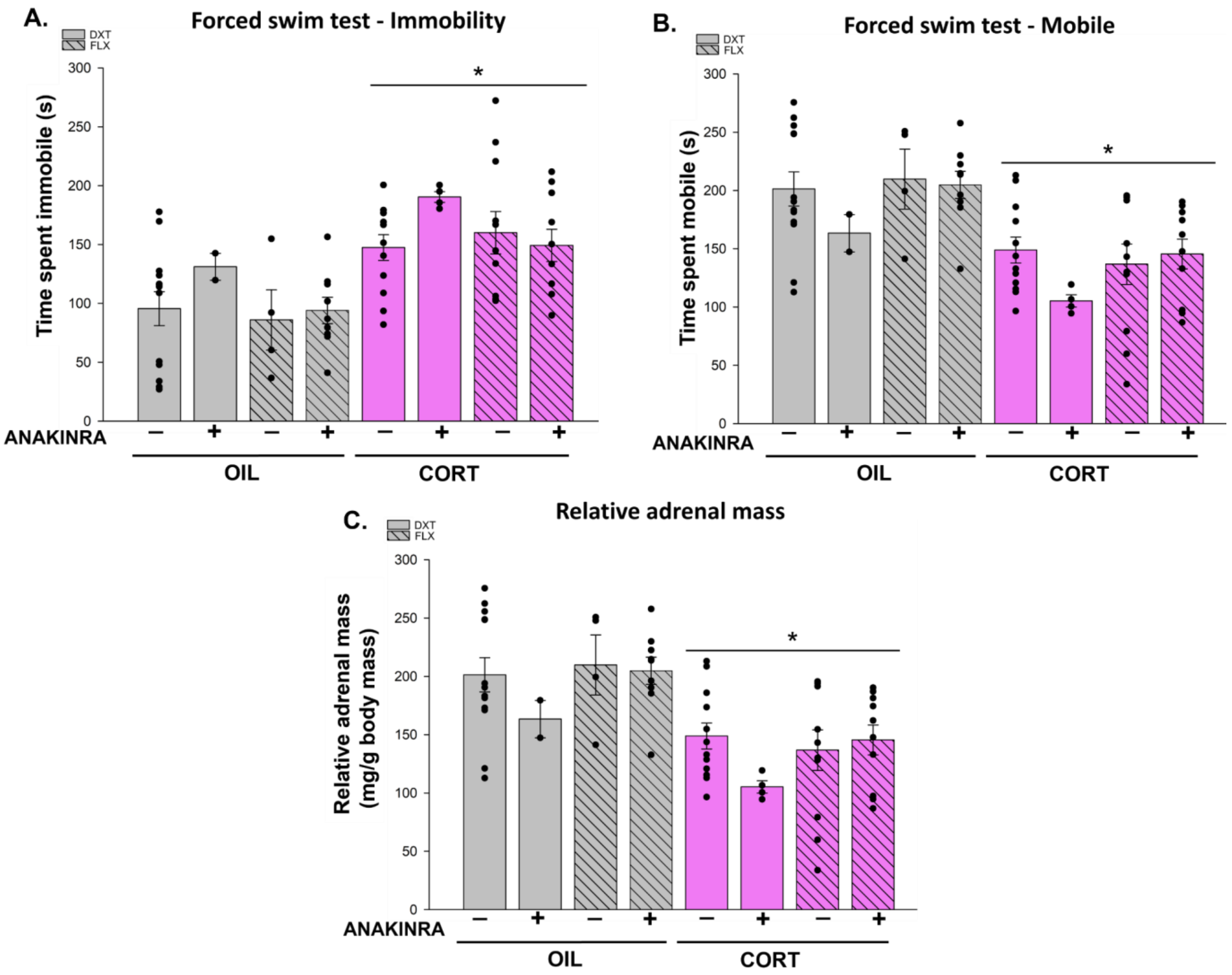


**Supplemental Figure 2. All treatment groups – Effects of postpartum corticosterone (CORT) and fluoxetine (FLX) treatment on immobility and mobility behaviour in the FST, and relative adrenal mass. A.** Postpartum CORT treatment significantly increased the time spent immobile in the FST. *Indicate significance at $p<0.05$. **B.** Postpartum CORT treatment significantly reduced time spent mobile in the FST. *Indicate significance at $p<0.05$. **C.** Postpartum CORT treatment significantly reduced relative adrenal mass. *Indicate significance at $p<0.05$. Data represented in means +standard error of the mean (SEM), overlaid with individual data points. n's = 2-13 per group.

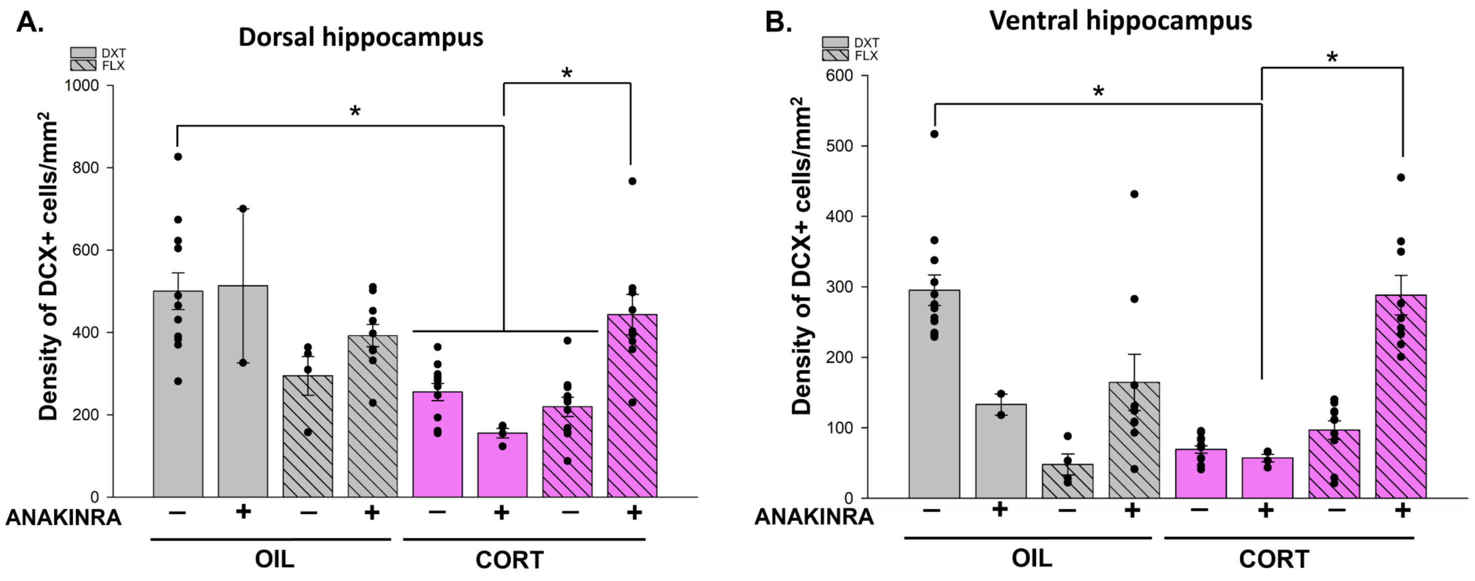


**Supplemental Figure 3. All treatment groups – Effects of postpartum corticosterone (CORT) and fluoxetine (FLX) treatment on dorsal and ventral DCX+ cells.**
**A.** Postpartum CORT significantly decreased the density of DCX+ cells in the dorsal hippocampus. FLX+KIN rescued these effects. *Indicate significance at $p<0.05$. **B.** Postpartum CORT significantly decreased the density of DCX+ cells in the ventral hippocampus. FLX+KIN rescued these effects. *Indicate significance at $p<0.05$. Data represented in means +standard error of the mean (SEM), overlaid with individual data points. n's = 2-13 per group.

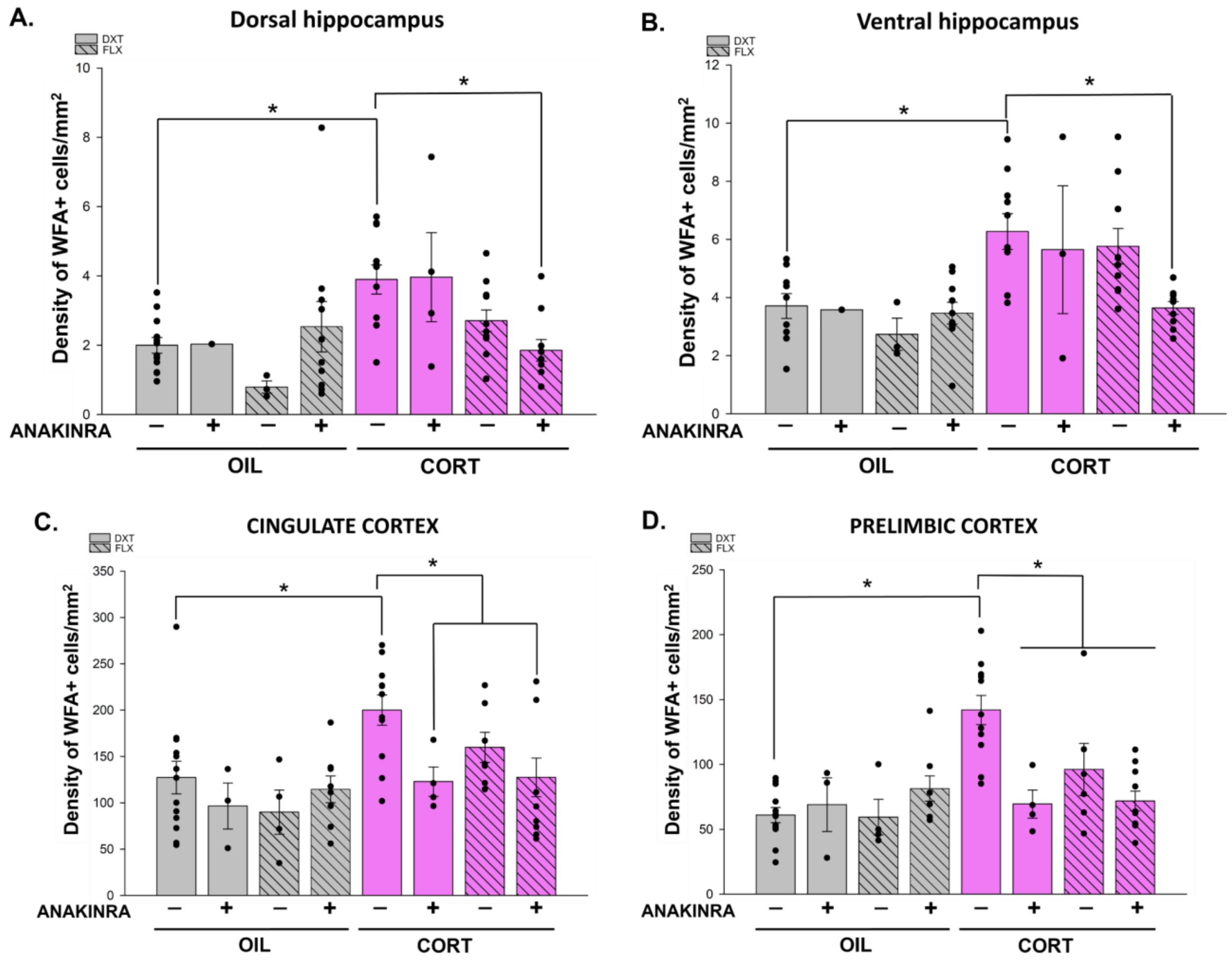


**Supplemental Figure 4. All treatment groups – Effects of postpartum corticosterone (CORT) and fluoxetine (FLX) treatment on cingulate and prelimbic cortices,and dorsal, ventral hippocampi WFA+ cells.**

**A.** Postpartum CORT significantly increased the density of WFA+ cells in the dorsal hippocampus. FLX+KIN rescued these effects. *Indicate significance at $p<0.05$. **B.** Postpartum CORT significantly increased the density of WFA+ cells in the ventral hippocampus. FLX+KIN rescued these effects. *Indicate significance at $p<0.05$. **C**. Postpartum CORT significantly increased the density of WFA+ cells in the cingulate cortex. Both KIN alone, and FLX+KIN together rescued these effects. Indicate significance at $p<0.05$. **D.** Postpartum CORT significantly increased the density of WFA+ cells in the prelimbic cortex. Both postpartum KIN alone, FLX alone and FLX+KIN together rescued these effects. Indicate significance at $p<0.05$. Data represented in means +standard error of the mean (SEM), overlaid with individual data points. n's = 3-13 per group.
